# Combinatorial adapter targeting enables AND-gate activation of AdCAR T cells in pancreatic cancer but reveals donor-dependent activation thresholds

**DOI:** 10.64898/2026.07.03.736407

**Authors:** Camille Dourlens, Kim Vanderliek, Olaf Hardt, Daniel Schäfer

**Author notes:** **Correspondence:** Daniel Schäfer.

## Abstract

Pancreatic ductal adenocarcinoma (PDAC) remains a lethal malignancy with limited therapeutic options, underscoring the need for innovative treatments. Chimeric antigen receptor (CAR) therapy has transformed hematologic malignancies but faces key challenges in solid tumors, particularly on-target/off-tumor toxicity and antigen heterogeneity. Adapter CAR (AdCAR) platforms offer enhanced control by decoupling antigen recognition from CAR activation, enabling controllable, reversible, and multi-antigen targeting. Recent studies suggest AdCARs can function as an AND-gate using combinations of adapter molecules at controlled surface densities. This defines activation thresholds, termed the Surface Activation Matrix, that restricts full activation to tumor cells overexpressing the target antigen combination, thereby reducing off-tumor toxicity. In this study, we evaluated its applicability to PDAC using adapters targeting CD318, TSPAN8 and CD66c. We systematically evaluated single and combinatorial adapter dosing in co-culture assays with AsPC1 cells, in a donor-dependent manner. Low concentrations of individual adapters were non-cytotoxic, whereas combining them at identical sub-threshold doses restored potent tumor killing, demonstrating that AdCAR activation depends on cumulative adapter density rather than total amount. However, the activation threshold required for AND-gate cytotoxicity varied between donors, highlighting the need for patient-specific titration to achieve selective tumor killing. These findings validate that AdCAR T cell activity in PDAC can be finely tuned through adapter concentration and combinatorial targeting, enabling selective tumor recognition while minimizing on-target/off-tumor toxicity. This flexible, safety-oriented strategy supports targeting heterogeneous PDAC tumors, though donor-dependent variability remains a critical consideration for clinical implementation.

## 1 Introduction

Pancreatic ductal adenocarcinoma (PDAC) remains a formidable challenge, as the third leading cause of cancer death in the U.S., causing ∼8% of deaths despite ∼3% of cases^1^. The ∼13% 5-year survival reflects limited curative options, mainly surgery, with ∼80% of cases unresectable at diagnosis. This underscores the need for new therapies^2,3,4,5^.

Chimeric Antigen Receptor (CAR) T cells, engineered to specifically recognize and eliminate target cells, have revolutionized adoptive immunotherapy and are now being evaluated as standard care for several hematological malignancies^6,7^. However, conventional CAR therapy faces is limited by single-antigen targeting and potential toxicity from uncontrolled activity or on-target/off-tumor effects^8^. To overcome these challenges, modular CAR platforms decouple antigen recognition from T cell activation via adapter or switch molecules^9^. These systems use a universal receptor that engages a soluble adapter binding the tumor antigen, enabling precise control and rapid retargeting without re-engineering the construct. Early examples include UniCAR^10^, switchable CAR (sCAR) with a peptide neo-epitope (PNE)^11^, and anti-FITC CARs recognizing fluorescein-labeled antibodies^12^. CD19-CARs have also been adapted using bridging proteins to redirect T cells toward alternative targets (e.g. HER2)^13^. These platforms provide temporal and dose-dependent control, as CAR T cells remain inert without the adapter, thereby improving safety and allowing fine-tuning^14^. Furthermore, shared adapter epitopes enable flexible targeting of multiple antigens, addressing antigen escape and tumor heterogeneity^15^. These advantages have lately driven increasing interest for the development of systems such as AdCAR^16^, SpyCatcher CAR^17^, SUPRA-CAR^18^, TRUE CAR^19^, and SNAP CAR^20^. Since 2020, six clinical trials have evaluated some of these platforms, underscoring their versatility and potential to overcome the limitations of conventional CAR therapy^9^.

Here, we use the previously described AdCAR platform, which employs an anti-biotin CAR recognizing a Linker-Label Epitope (LLE) generated via specific biotinylation chemistry^16^. This enables biotin-tagged adapters to bridge CAR T cells to diverse tumor antigens, allowing controlled activation and combinatorial targeting. Recently, CD318, TSPAN8 and CD66c have been identified as promising targets overexpressed in PDAC^21^, but their low basal expression in healthy tissues raises toxicity concerns. To further minimize this risk, an additional control layer is introduced: AdCAR activation requires a threshold density of adapter molecules on the target cell surface. This threshold can be reached either by increasing the amount of a single-adapter or by combining multiple adapters targeting different antigens, even at low individual levels. This Surface Activation Matrix (SAM) strategy enables selective targeting of tumor cells^16^. In this “AND-gate” configuration, only cells expressing all targeted antigens reach the activation threshold, whereas healthy cells with lower or partial expression remain largely unaffected.

In this study, we applied this targeting strategy to CD318, TSPAN8 and CD66c^11^. To this end, biotinylated adapters against each antigen were used to systematically evaluate individual and combined dosing. We found that combining three otherwise inactive single-doses was sufficient to induce strong, specific anti-tumor activity. However, donor-specific variability remains a challenge for broader application.

## 2 Methods

### Bioinformatical data analysis

#### GEPIA2

The expression boxplots were generated using GEPIA2^22^. The dataset used was PAAD, with matched TCGA normal and GTEx data. The method for differential analysis was one-way ANOVA, using disease state (Tumor or Normal) as variable for calculating differential expressions. The expression data was first log_2_(TPM+1) transformed for differential analysis and the log_2_FC was defined as median(Tumor) - median(Normal). Genes with higher |log_2_FC| values and lower q values than pre-set thresholds are considered differentially expressed genes. The complete data is at http://gepia2.cancer-pku.cn/#analysis.

#### EMBL-EBI, Expression Atlas

The heatmap was generated on *Homo Sapiens* set. Data is from the genotype-tissue expression (GTEx) project v8 from RNA-Seq data^23^ and available at https://www.ebi.ac.uk/gxa/experiments/E-GTEX-8/.

### Cell lines and culturing conditions

AsPC1 and SupT1 (ATCC) cell lines were cultured in RPMI media (Biowest), supplemented with 2mM L-glutamine (Biowest) and 10% Fecal Calf Serum (FCS) (Eximus). HEK293T (ATCC) cell line was cultured in DMEM High-Glucose media (Biowest), supplemented with 10% FCS.

### Adapter molecule conjugation

Pure antibodies were obtained from Miltenyi Biotec and rebuffered at 1 mg in a 0.1 M NaHCO3 solution (Sigma-Aldrich) via a NAP-5 column (Cytiva). Antibody modification was performed at room temperature for 1h in NaHCO3 buffer using a 2-fold molar excess of biotin-LC-LC-NHS (ThermoFisher Scientific), previously dissolved at 10 mg/mL in anhydrous DMSO (ThermoFisher Scientific). The reaction mix was purified using NAP columns and samples were rebuffered in PBS (VWR) for storage. Concentration was measured by absorption at 280 nm with a Nanodrop (ThermoFisher Scientific). To validate the successful biotinylation, a flow cytometry assay was performed on AsPC1 cells using anti-biotin-PE as secondary-staining.

### Generation of AdCAR-construct and lentivector particles

The AdCAR used in this study was produced as previously described^16^. Lentiviral (LV) particles were generated by the transient transfection of HEK293T cells using polyethylenimine. After 48h, supernatant containing lentiviral pseudo-particles were concentrated by overnight centrifugation and resuspended in TexMACS™ Medium (Miltenyi Biotec) before storing at −70 °C. LV titers were determined by the transduction of SupT1.

### Isolation of T cells and generation of AdCAR T cells

Peripheral blood mononuclear cells (PBMCs) were isolated by PanColl (PanBiotech) density gradient centrifugation of whole blood. T cells were purified using the human Pan T Cell Isolation Kit and activated in TexMACS™ Medium supplemented with human T Cell TransAct™ and 12.5 ng/mL of both recombinant human interleukins IL-7 and IL-15 (all Miltenyi Biotec). 24 hours after activation, T cells were transduced with LV. Three days post activation, TransAct™ was washed out and T cells were cultured with TexMACS™ Medium supplemented with IL-7 and IL-15. AdCAR transduction efficiency was assessed by flow cytometry.

### Cytotoxicity assays and live-imaging

Target cells were inoculated at densities of 2E4 cells per well with AdCAR^+^ T cells in an effector-to-target (E:T) ratio of 5:1. Sterile biotinylated adapters were added to each well at the desired final concentration. Cytotoxicity was measured as a decrease in green surface area with an IncuCyte® S3 Live-Cell Analysis System (Sartorius) and the supplied software (v2019A). The green surface area was normalized to the first measurement. For heatmap generation, slopes were calculated using the SLOPE function on Microsoft Excel from Incucyte measurements. Negative slopes between −3 and −0.005 (green) indicated tumor cell killing, whereas slopes between −0.005 and 0.001 (orange) indicated growth inhibition. Positive slopes >0.001 (red) indicated tumor cell growth. Endpoint cytotoxicity was calculated as the percentage decrease or increase of green surface area from the first to the last measurement. At the end of the assay, T cells were later analyzed for their activation markers and 100 μl of medium was used for cytokine release using a human MACSplex cytokine 12 kit (Miltenyi Biotec). Cytokine data were analyzed using MACSPlexInspectoR software (v1.0) from Miltenyi Biotec (https://www.miltenyibiotec.com/DE-en/resources/tools/macsplex-inspector.html). Assay workflow is presented in supplementary figure S1.

### Flow cytometry

All samples were measured on MACSQuant® Analyzer 10 or 16 from Miltenyi Biotec. Samples were analyzed using the MACSQuantify™ Software (Miltenyi Biotec, v2.13.0), FlowJo (BD Biosciences, v10.10.0), GraphPad Prism (GraphPad Software, v10.1.2) and Microsoft Excel for Microsoft 365 MSO (Microsoft, v2502). Staining conducted with antibody conjugates from Miltenyi Biotec was performed as recommended by the supplier. Dead cells were excluded using 7-AAD. All antibodies are listed in supplementary table 1. Gating strategy for tumor and CAR T cell phenotyping after co-cultures is shown in supplementary figure S2.

### Statistics

Graphs and statistical analysis were generated using GraphPad Prism 10. Killing curves are presented as mean of technical replicates (n = 2) per condition. Error bars and data in barplots are presented as means ± SD or means from the different replicates, as indicated in figure legends. Statistical significance was assessed using a One-Way ANOVA test.

## 3 Results

### Selection of CD318, TSPAN8 and CD66c as suitable PDAC antigen targets for combination

To implement the combinatorial AdCAR strategy (figure 1.A), we selected three relevant PDAC-associated antigens: CD318 *(CDCP1)*, CD66c *(CEACAM6)* and TSPAN8, previously shown to be overexpressed in PDAC^21^. Gene expression analysis revealed elevated levels of all three targets in primary pancreatic tumors compared to normal tissues, with CD66c displaying the highest differential expression (figure 1.B). Expression on AsPC1 cells confirmed high levels of CD318 (83.6%) and TSPAN8 (85.3%), and near-uniform CD66c expression (99.2%) (figure 1.C), validating AsPC1 as a suitable model. As these antigens are not only strongly expressed in PDAC but exhibit also basal expression in some normal tissues, an AND-gate targeting strategy appears to be well suited for their application (figure 1.D). These patterns suggest that single-CAR targeting may therefore carry on-target/off-tumor toxicity risk. However, the lack of major overlapping expression in healthy tissues makes these antigens suitable for combinatorial AdCAR targeting, as simultaneous overexpression occurs primarily in PDAC. Furthermore, these antigens have previously demonstrated good compatibility with CAR systems, further supporting their candidacy for safe and effective combinatorial targeting in PDAC^21,24^.

**Figure 1.**
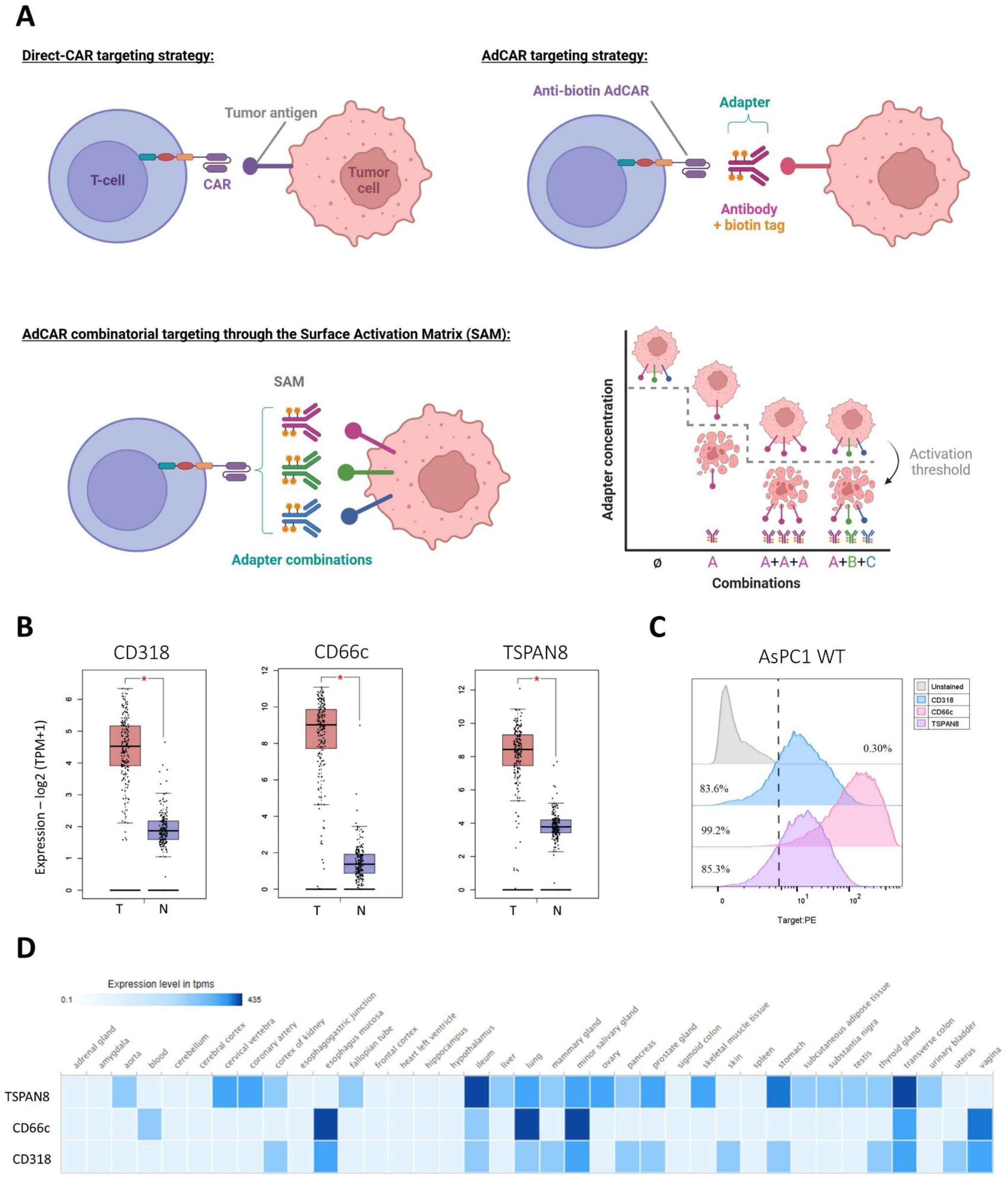
AdCAR T cell targeting strategy and CD318, CD66c and TSPAN8 expression in pancreatic adenocarcinoma versus healthy tissues. **A)** Comparison between mono-CAR and AdCAR targeting strategies. Direct CARs bind tumor cell antigens via scFvs. AdCARs separate antigen recognition from CAR activation, triggering T cell activation only in the presence of specific adapter molecules (e.g., biotinylated antibodies). In the surface activation matrix concept (SAM), AdCAR activation requires a threshold density of adapter molecules on the target surface, achieved either by high levels of a single-adapter (A+A+A) or by combining distinct sub-threshold adapters (A+B+C). Adapter combinations enable full tumor killing without significant cytotoxicity from single targeting. Adapted from Seitz et al. 2021^16^ and created with BioRender.com. **B)** Barplots of CD318, CD66c and TSPAN8 gene expression in healthy pancreas (N=171) versus pancreatic adeno-carcinoma tumor (T=179). Data were retrieved as log₂(TPM+1) with an absolute log₂ fold-change (log₂FC) cutoff value of 1 and a p-value cutoff of 0.01. Statistical significance was assessed using one-way ANOVA. Graphs were extracted from GEPIA2^22^. **C)** Histogram showing flow cytometry staining of AsPC1 tumor cells showing the expression of CD318, TSPAN8 and CD66c compared with unstained control. Percentages indicate the positive fractions determined using the unstained control for gating. Data is representative of one sample from a technical triplicate. **D)** Heatmap of TSPAN8, CD66c and CD318 gene expressions (in transcript per millions) across healthy tissues from GTEx project v8 data. Graphs were extracted from EMBL-EBI/Expression Atlas^23^.

### Biotinylated adapters enable efficient target binding and concentration-dependent cytotoxicity

CD318, TSPAN8 and CD66c antibodies were biotinylated to generate the adapter molecules. Successful biotinylation was confirmed by flow cytometry using adapter concentrations ranging from 1 to 100 ng/mL. A clear signal separation from unstained controls was observed at 100 ng/mL (figure 2.A), which progressively decreased at lower concentrations. CD66c adapters showed the strongest binding, consistent with high CD66c expression on AsPC1 cells, with distinct signal even at 1 ng/mL. In contrast, CD318 and TSPAN8 exhibited weaker staining, in line with their lower antigen expression.

**Figure 2.**
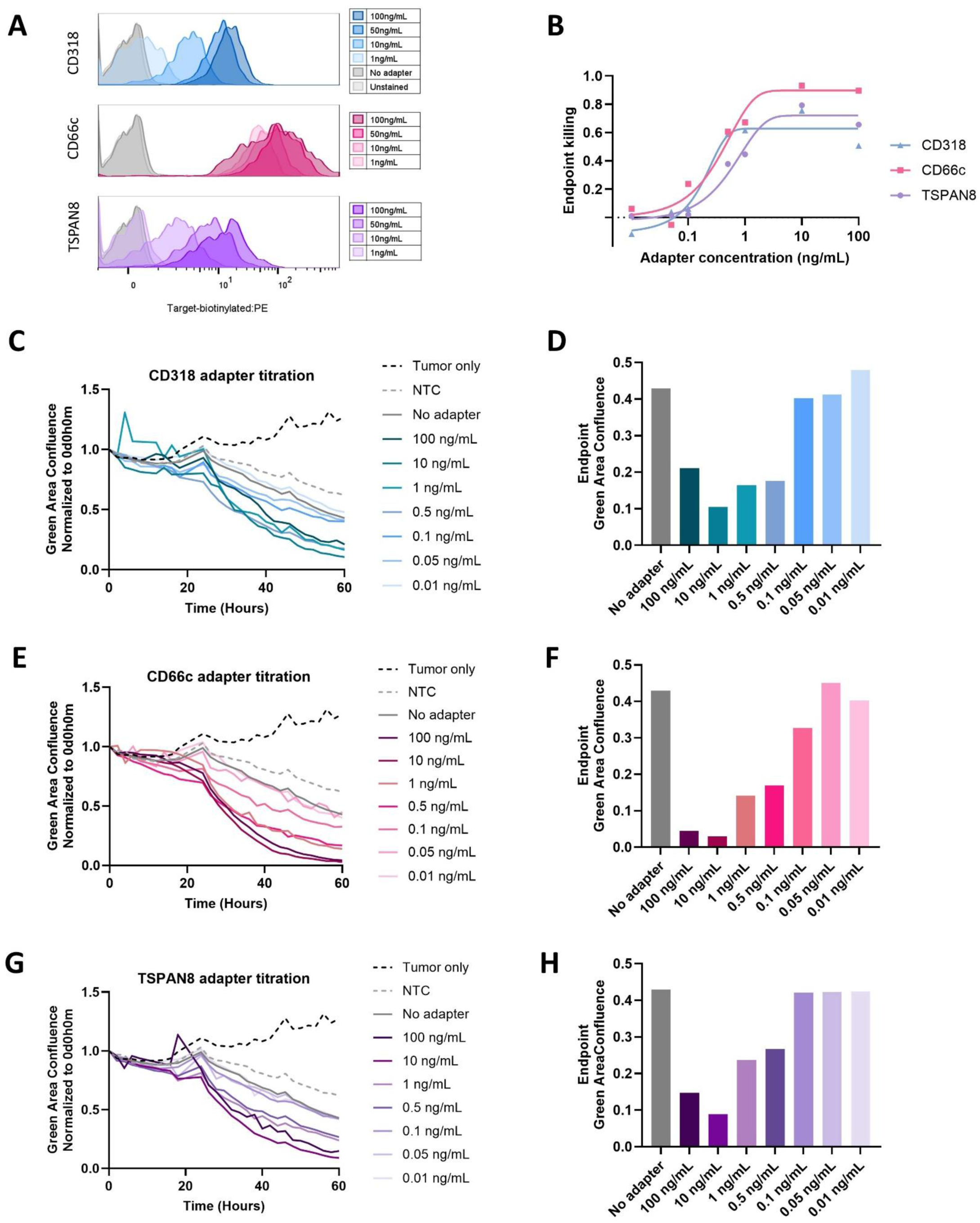
Titration and cytotoxic characterization of individual biotinylated AdCAR adapters against AsPC1 pancreatic cancer cells. **A)** Histogram showing flow cytometry staining with biotinylated adapters as primary staining and subsequent anti-biotin-PE staining to confirm successful biotinylation. AsPC1 tumor cells were incubated with varying concentrations (1 to 100 ng/mL) of biotinylated CD318, TSPAN8 or CD66c compared with unstained controls and samples stained without adapter molecules. Data is representative of one sample from a technical duplicate. **B)** Comparison of endpoint cytotoxicity after a 60 h killing assay using individual CD318, CD66c and TSPAN8 adapters at concentrations ranging from 0.01 to 100 ng/mL. Data were fitted using a nonlinear sigmoidal dose-response model using GraphPad Prism. **C)E)G)** Representative killing curves showing GFP-positive tumor cell confluence (%) normalized to the first time point for AsPC1 cells co-cultured with AdCAR T cells (E:T ratio of 5:1) in the presence of increasing concentrations (0.01-100 ng/mL) of the CD318 **C)**, CD66c **E)**, or TSPAN8 **G)** adapter. **D)F)H)** Endpoint cytotoxicity after 60 h for the CD318 **D)**, CD66c **F)**, and TSPAN8 **H)** adapters across the indicated concentration range. **B-H)** Data points represent the mean of three technical replicates of one donor for each condition.

We next evaluated cytotoxic activity across a concentration range (0.01-100 ng/mL) in 60 h killing assays. Consistent with expression levels, CD66c adapters showed the highest cytotoxic potency, reaching a plateau of ∼93% tumor clearance at the experiment endpoint with 10 ng/mL, whereas CD318 and TSPAN8 achieved ∼75% and ∼79% clearance, respectively (figure 2.B). All adapters displayed clear dose-dependent killing (figures 2.C to 2.H). Concentrations below 0.5 ng/mL induced minimal cytotoxicity, comparable to non-transduced T cell controls, indicating insufficient adapter availability for productive activation. The 0.5-10 ng/mL range corresponded to the inflection region of the sigmoidal response curve and therefore represents the most informative concentration window for subsequent AND-gate studies (figure 2.B).

Specificity of the AdCAR system was further assessed using saturating adapter concentrations (1000 ng/mL; supplementary figure S3). At this concentration, AdCAR T cells combined with CD318, TSPAN8 or CD66c adapters induced strong cytotoxicity after 70 h, whereas neither AdCAR T cells nor adapters alone exhibited any cytotoxic activity, similar to non-transduced T cells and tumor-only controls. In addition, AdCAR T cells combined with a biotinylated CD19 isotype control adapter showed no activity against CD19-negative AsPC1 cells, confirming system specificity. Together, these data define the functional characteristics of the biotinylated adapters, demonstrate the specificity of the AdCAR system, and establish an optimal concentration range for subsequent combinatorial AND-gate experiments.

### Combinatorial adapter targeting enables AdCAR cytotoxicity while single-dosing fails

Adapters were tested individually and in combination at varying concentrations using AdCAR T cells co-cultured with AsPC1 cells at E:T ratio of 5:1. Successful AdCAR expression was confirmed (supplementary figure S4). No cytotoxicity was observed in non-transduced (NTC) controls. Moreover, AdCAR induced-cytotoxicity occurred exclusively in the presence of the antigen-specific conjugated adapters (figure 3.A). Cytotoxicity was observed with single-adapters at ≥20 ng/mL and increased in a dose-dependent manner (figure 3.B). Combinations of two or three adapters consistently enhanced killing compared to single-adapters (figures 3.A and 3.C). Above 5 ng/mL per adapter, all conditions induced cytotoxicity, indicating sufficient activation with single targeting. However, to establish AND-gate behavior, we sought a sub-threshold concentration at which single-adapters and dual combinations were insufficient to induce killing, while triple-adapter engagement resulted in cytotoxicity. Due to higher CD66c intrinsic activity, this condition was achieved only at 1 ng/mL. At this level, single-CD318 and TSPAN8 adapters permitted tumor growth, while single-CD66c and dual-adapter combinations inhibited growth without killing. In contrast, only the triple-adapter combination induced robust cytotoxicity.

**Figure 3.**
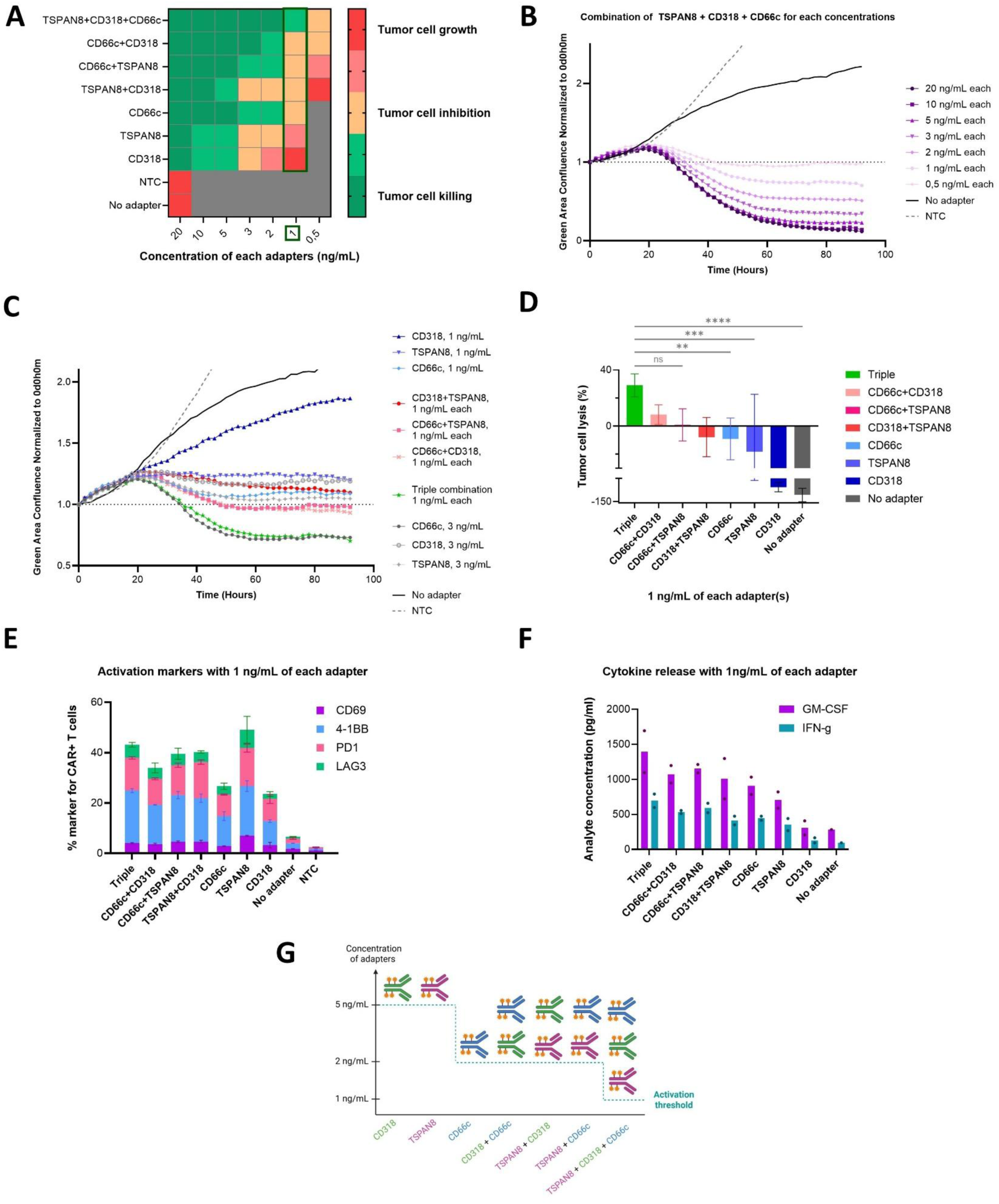
In vitro evaluation of combinatorial AdCAR targeting under different adapter conditions. **A)** Heatmap summarizing AdCAR cytotoxicity on AsPC1 at E:T ratio of 5:1, across adapter combinations and concentrations (in ng/mL), acquired by the Incucyte system. Colors were assigned based on the slope values of the killing curves: green indicated cytotoxicity, orange indicated tumor cell growth inhibition and red indicated tumor cell growth. Each condition shows mean of technical replicates from one donor. **B)** Killing curves showing GFP-based tumor cell confluence normalized to the first time point of the triple-adapter combination (TSPAN8+CD318+CD66c) tested from 20 to 0.5 ng/mL versus no-adapter control, on AsPC1 E:T ratio of 5:1. Each data point represents the mean of technical duplicates per condition. **C)** Killing curves comparing different adapter combinations at 1 ng/mL each, versus no-adapter and single-adapter controls at 3 ng/mL, on AsPC1 E:T 5:1. Each data point represents the mean of technical duplicates per condition. **D)** Barplot of endpoint cytotoxicity for each adapter combination at 1 ng/mL per adapter. Mean ± SD of two technical duplicates from 4 images per well. Statistical significance was assessed using a One-Way ANOVA test; non-significant (ns), p < 0.0001(****). **E)** Activation markers CD69, 41BB, PD1 and LAG3 and **F)** cytokine levels (GM-CSF and IFN-g) measured at assay endpoint for each adapter combination at 1 ng/mL. Each data point represents the mean ± SD of technical duplicates per condition. **G)** Summary of donor-specific activation thresholds for different adapter combinations. Created with BioRender.com.

Detailed GFP-based killing kinetics at 1 ng/mL showed sustained tumor reduction only in the triple condition, with 29% elimination at endpoint versus 8% for CD66c+CD318 and none for others (figure 3.D). Notably, the combination of three adapters at 1 ng/mL each (so with a total of 3 ng/mL of combined adapters) mediated more effective killing than individual adapters used at 3 ng/mL each (except for CD66c), demonstrating that AdCAR activation depends on cumulative adapter density rather than total adapter amount (figure 3.C). Together, these results identify 1 ng/mL as the optimal concentration for enforcing AND-gated AdCAR activation for this donor and highlight CD66c as the dominant contributor to cytotoxicity within adapter combinations.

The activation-marker profiles were broadly similar across conditions, with lower expression for single-CD318 and CD66c (figure 3.E). Lower activation for CD318 adapter alone is consistent with the reduced cytotoxicity observed before. AdCAR T cells got activated exclusively in the presence of adapters. Despite the triple-adapter combination eliciting significantly higher tumor cell killing, there was no proportional increase in activation-marker expression compared with other combinations. Specifically, after a 90-hour assay, CD69 remained low across all conditions, indicating sustained rather than early T cell activation. 4-1BB expression was also comparable, suggesting consistent co-stimulatory signaling. PD-1 expression was lower than CD69 and 4-1BB but followed similar trends across conditions. LAG-3 expression remained uniformly low, indicating minimal T cell exhaustion, except for the condition with TSPAN8 adapter only.

Cytokine secretion mirrored these trends: GM-CSF and IFN-γ were slightly elevated in the triple-adapter condition and reduced for single-CD318, while most other cytokines were undetectable (figure 3.F). The presence of elevated GM-CSF and IFN-γ indicates antigen-driven CAR T cell activation, with IFN-γ reflecting cytotoxic effector function and GM-CSF suggesting engagement of inflammatory and myeloid-activating pathways. However, the absence of IL-2 and TNF-α secretion was detected, maybe due to limited CAR T cell activation and proliferative capacity (supplementary figure S5). IFN-α, IL-4, IL-6, IL-10, IL-12, and IL-17A were not detectable, indicating a low risk of pronounced inflammatory, regulatory, or alternative T helper polarization signals. Interestingly, low levels of IL-5 and IL-9 were observed across all conditions except CD318, revealing a modest induction of non-classical helper T-cell-associated cytokines. Without antigen-specific adapters, cytokine production remained very low, which, together with the absence of activation, confirms the lack of AdCAR activity in the absence of adapter.

Collectively, these results indicate that the enhanced cytotoxicity observed with triple-adapter engagement arises from cumulative antigen recognition rather than overt T cell hyperactivation, supporting the concept that AND-gate AdCAR T cells can achieve potent, target-restricted killing while maintaining controlled and sustained activation without inducing T cell exhaustion.

### AND-gate AdCAR cytotoxicity requires donor-specific adapter titration

We first demonstrated that an AND-gate activation profile could be achieved by carefully selecting the concentration range of the three-adapter combination to reach the killing threshold. Selective cytotoxicity was observed exclusively with the triple-adapter combination at an activation threshold of 1 ng/mL per adapter (figure 3.G). We next tested whether AND-gate activation generalizes across donors.

A second donor showed similar trends, with increased killing at higher concentrations and enhanced cytotoxicity with combined adapters (figures 4.A and 4.B). However, the activation threshold shifted: selective killing via triple adapters occurred between 5 and 10 ng/mL, compared to 1 ng/mL for the first donor. At 10 ng/mL, both triple and TSPAN8+CD66c combinations induced cytotoxicity, while single adapters remained inactive, preserving AND-gate behavior although with a requirement for only two adapters rather than three. Importantly, this dual-adapter AND-gate configuration is still expected to reduce off-tumor toxicity, as TSPAN8 and CD66c exhibit largely non-overlapping expression profiles in healthy tissues (figure 1.D). In contrast, a comparable activation threshold involving CD318 and TSPAN8 would pose higher risk, given their overlapping expression in healthy colon and, to a lesser extent, stomach tissues.

**Figure 4.**
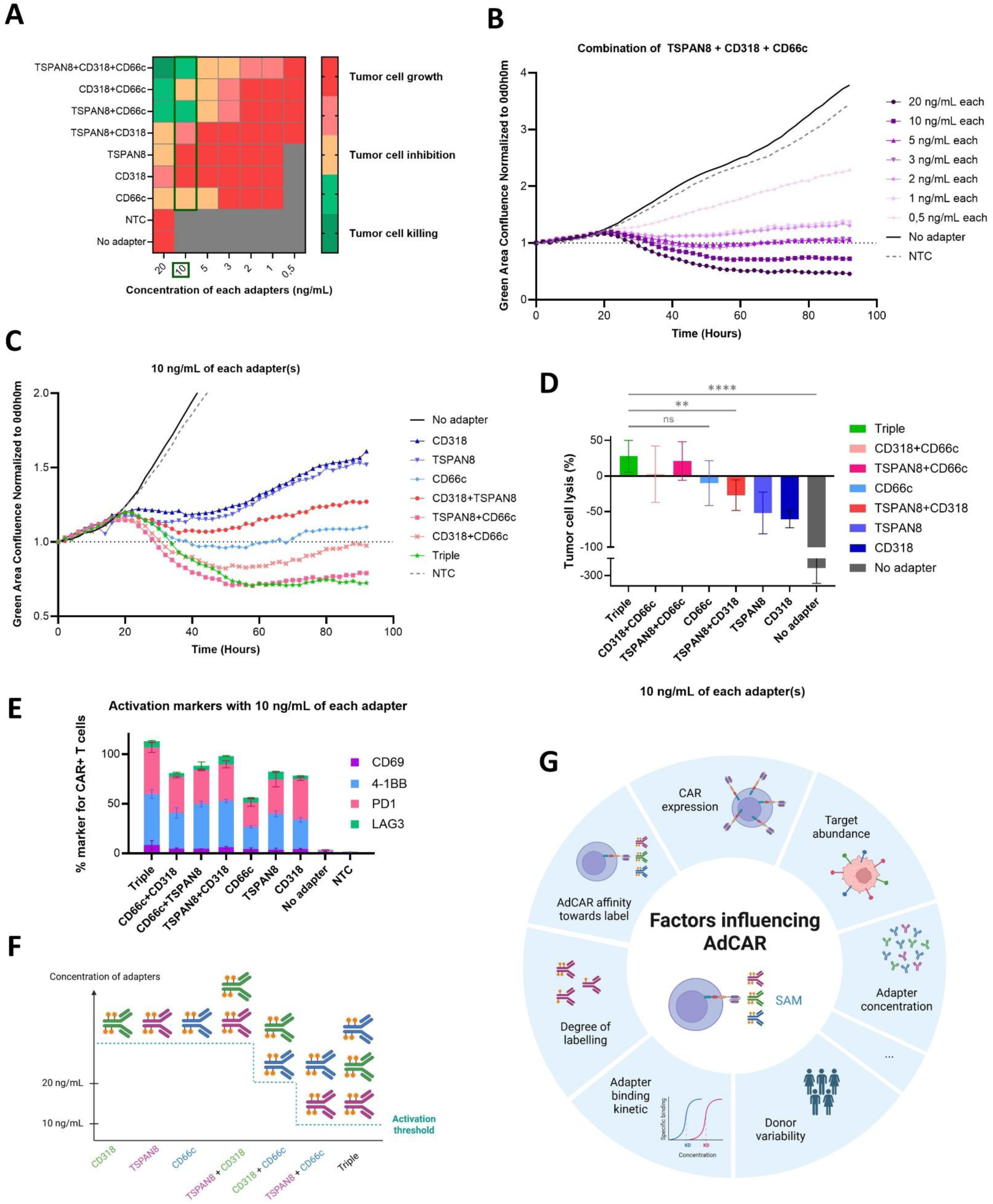
In vitro evaluation of donor-variability in the combinatorial AdCAR targeting system with a second donor. **A)** Heatmap summarizing AdCAR cytotoxicity on AsPC1 at E:T ratio of 5:1 in a second donor, across adapter combinations and concentrations (in ng/mL), acquired by the Incucyte system. Colors were assigned based on the slope values of the killing curves: green indicated cytotoxicity, orange indicated tumor cell growth inhibition and red indicated tumor cell growth. Each condition shows mean of technical replicates. **B)** Killing curves showing GFP-based tumor cell confluence normalized to the first time point of the triple-adapter combination (TSPAN8+CD318+CD66c) tested from 20 to 0.5 ng/mL versus no-adapter control, on AsPC1 E:T ratio of 5:1. Each data point represents the mean of technical duplicates per condition. **C)** Killing curves comparing different adapter combinations at 10 ng/mL each, versus no-adapter, on AsPC1 E:T 5:1. Each data point represents the mean of technical duplicates per condition. **D)** Barplot of endpoint cytotoxicity for each adapter combination at 10 ng/mL per adapter. Mean ± SD of two technical duplicates from 4 images per well. Statistical significance was assessed using a One-Way ANOVA test; non-significant (ns), p < 0.0001(****). **E)** Activation markers CD69, 41BB, PD1 and LAG3 measured at assay endpoint for each adapter combination at 10 ng/mL. Each data point represents the mean ± SD of technical duplicates per condition. **F)** Summary of the second donor-specific activation thresholds for different adapter combinations. Created with BioRender.com. **G)** List of factors that may influence AdCAR responses and SAM kinetics and could potentially account for the observed variability. Created with BioRender.com

A striking observation was the substantially lower activation threshold in the first donor compared to the second (∼10 ng/mL versus 1 ng/mL per adapter), representing a 10-fold difference. At 10 ng/mL, cytotoxicity appeared less clearly stratified than in the first donor, although enhanced killing remained evident with the triple-adapter and TSPAN8+CD66c combination (figures 4.C and 4.D). Consistent with the first donor, CD66c adapter displayed relatively higher intrinsic activity than TSPAN8 and particularly CD318, likely explaining the increased cytotoxicity in CD66c-containing combinations. Likewise, 4-1BB and PD-1 expression was elevated in the triple combination, reduced with CD66c alone, and comparable across other conditions (figure 4.E).

Collectively, these data demonstrate that the adapter concentration required for AND-gate AdCAR activation is strongly donor-dependent, varying up to 10-fold for these two examples (figure 4.F). This donor variability suggests that adapter dosing must be individually optimized to achieve selective tumor targeting while minimizing on-target/off-target toxicity. We propose several factors that may influence AdCAR performance and the SAM threshold, including CAR expression levels, target antigen abundance, adapter concentration, donor-to-donor variability, adapter binding kinetics, and degree of labeling, as well as AdCAR affinity for the labeling moiety and additional yet unidentified parameters (figure 4.G). Accordingly, while the AND-gate AdCAR system represents a powerful strategy to enhance specificity, its clinical implementation may therefore require patient-specific titration and tight control of all these parameters, adding a layer of complexity for translational applications.

## 4 Discussion

The remarkable success of CAR T cell therapies in B cell malignancies has not been yet fully extended to other cancers. This is primarily due to the presence of distinct lineage markers in B cell malignancies such as CD19^25^, CD20^26^, CD22^27^, and BCMA^28^, which enable precise targeting without harming essential tissues. In contrast, solid tumor antigens are often expressed in healthy tissues, increasing the risk of toxicity. Consequently, CAR T cell clinical application in solid tumors is limited by two major challenges: on-target/off-tumor toxicity and antigen heterogeneity^8,29^. Many candidate targets investigated in clinical studies are expressed not only on tumor cells but also on healthy tissues, which has already resulted in severe and sometimes fatal side effects^30,31,32^. Additionally, solid tumor antigens are typically expressed at lower and more heterogeneous levels than B cell lineage markers, creating a permissive environment for clonal evolution, often culminating in the emergence of escape variants that bypass CAR-T-mediated immunosurveillance^33^. In PDAC, three recently identified targets (CD318, CD66c, TSPAN8) illustrate these challenges^21^. While CD318 appears to have a relatively favorable expression profile, CD66c and TSPAN8 are also expressed in several epithelial tissues, underscoring the risk of on-target/off-tumor toxicity. To address these limitations, we sought to extend an existing modular CAR approach by incorporating these three newly identified targets, with the aim of improving both safety and targeting precision in PDAC.

Modular CAR platforms have demonstrated encouraging efficacy in preclinical and early clinical studies, highlighting their potential as adaptable and controllable immunotherapeutic strategies^9^. For example, the ongoing Phase I trial of UniCAR02-T-CD123 (NCT04230265) in hematologic malignancies illustrates the clinical feasibility of such systems. However, some modular CAR designs may induce immunogenicity, particularly those incorporating non-human components such as avidin, anti-FITC CARs, or PNE-based systems^11,12,34^. In contrast, the AdCAR platform redirects T cells via an LLE-specific CAR, like a biotin-CAR, which are expected to be less immunogenic, and therefore more suitable for clinical applications. Moreover, AdCAR enables controlled targeting of different antigens via exchangeable biotinylated antibody adapters, compatible with any FDA- or EMA-approved antibody. This allows sequential or simultaneously targeting against multiple tumor antigens, increasing flexibility and improving the ability to address antigen escape and heterogeneity ^35^.

Although combinatorial targeting can mitigate antigen escape, it does not inherently resolve on-target/off-tumor toxicity. To address this, we further applied the SAM strategy, introducing an AND-gate mechanism for AdCAR activation^16^. Here, multiple adapters are required to trigger cytotoxicity, enabling precise control of T cell activity. Previous studies have shown that combinatorial targeting involving up to five targets can achieve selective tumor killing while minimizing on-target/off-tumor effects. In this work, we investigated whether similar AND-gate activity could be extended to PDAC using a reduced number of targets (here, three).

Our results demonstrate that AdCAR T cells can be precisely tuned through combinatorial targeting of three PDAC-associated antigens (CD318, TSPAN8, CD66c) achieving selective tumor killing while minimizing off-tumor toxicity. Single-adapters at low concentrations were insufficient to induce cytotoxicity, whereas combining them at sub-threshold doses restored potent killing. This confirms that AdCAR activation depends on cumulative adapter density rather than total adapter amount, enabling a functional AND-gate mechanism. Notably, triple-adapter combinations at low concentrations induced stronger cytotoxicity than higher single-adapter doses despite lower total load. Importantly, enhanced cytotoxicity in the triple-adapter condition was not associated with excessive T cell activation or early exhaustion. Activation markers and cytokine profiles remained controlled, with low CD69 and modest changes in 4-1BB, PD-1 and LAG-3. Cytokine release followed similar patterns, supporting cooperative antigen recognition rather than hyperactivation. This controlled response may improve safety and T cell persistence^14,36^. We observed a remarkable difference in the cytokine profile of the first donor compared with what is typically expected for conventional CAR T cells, particularly the absence of detectable IL-2 and TNF-α production. Previous studies have reported that AdCAR T cells can exhibit cytokine secretion patterns distinct from those of canonical CAR T cells at early timepoints^37^. However, the extent to which these reported differences account for the observations made in our study remains unclear.

A key limitation highlighted for this approach is the donor-dependent variability. While one donor achieved optimal AND-gate activity at low adapter concentrations, another required higher doses to reach comparable cytotoxicity. Between two donors, this 10-fold difference underscores the complexity of defining a universal dosing strategy and suggests that patient-specific tuning may be required. In one hand, reducing the number of adapters from five to three simplifies the system, enhances feasibility for clinical use, and broadens applicability to other cancer types, since fewer tumor-specific targets need to be identified. This represents a major practical advantage given the challenges of target discovery in oncology. On the other hand, donor-dependent activation thresholds suggest that a one-size-fits-all dosing strategy may not be sufficient, and additional targets or fine-tuning of adapter concentrations may be required to ensure efficacy across a diverse patient population. As we also have seen tight therapeutic windows with the combination of three targets, the SAM approach would certainly profit from using even more targets. In PDAC, additional antigens such as CLA^21^, ROR1^38^, Claudin 18.2^39^, HER2^40^ or MSLN^41^ could be considered to increase flexibility and refine threshold control, but this introduces a trade-off between system simplicity and tunability.

Several factors may influence AdCAR performance and the SAM threshold, including CAR expression levels, target antigen abundance, adapter concentration, donor-to-donor variability, adapter-binding kinetics, and degree of labeling, as well as AdCAR affinity for the labeling moiety and additional yet unidentified parameters. Adapter concentration is unlikely to account for the observed differences, as this parameter was carefully controlled throughout the experiments. We also evaluated target antigen abundance and adapter binding characteristics, which showed only minor differences between adapters, with CD66c exhibiting the highest binding and expression levels. However, because identical target cells and the same adapter preparations were used across all donors, these factors are unlikely to explain the observed inter-donor variability. Nevertheless, they should be carefully considered when comparing different adapter production batches or when extending this approach to additional targets or tumor cell lines.

Differences in AdCAR expression can influence the overall AdCAR response, as normalization of the E:T ratio does not account for CAR density per cell, which influences functional avidity. Avidity reflects the combined strength of multiple receptor-ligand interactions and is influenced by CAR expression, signaling strength, and tumor antigen density^42^. It has been shown that tuning CAR affinity can impact T cell expansion and persistence, as demonstrated with low-affinity CD19 CARs^43^. In addition, T cell function is influenced by donor-dependent factors such as subset composition, differentiation state, transduction efficiency, tonic signaling, and overall cellular fitness and exhaustion^44,45^. These parameters are particularly relevant in the context of combinatorial targeting, where cooperative antigen recognition can enhance functional avidity and improve tumor killing in heterogeneous environments. However, although the percentage of CAR-positive cells in the second donor was approximately half that observed in the first donor, the MFI was comparable. This suggests that the observed differences are more likely attributable to donor-dependent factors rather than variations in transduction efficiency, particularly since all experiments were performed at comparable E:T ratios.

The degree of biotinylation was not assessed in this study. We do not expect this to account for the observed donor-to-donor variability, as identical adapter preparations were used across all experiments. However, differences in biotinylation between individual antibodies may contribute to the differential activity observed across targets. In particular, the stronger activity of the CD66c adapter may partially reflect biotinylation efficiency, although it is more likely driven by higher target expression levels on tumor cells. In this study, we primarily aimed to highlight the existence of inter-donor variability rather than to define the full range of activation thresholds in a statistically powered donor cohort. For broader generalization, future studies should systematically evaluate larger donor numbers together with adapter biotinylation degree, pharmacokinetics, and CAR density per cell to better define activation thresholds across donors and adapter lots. This would also facilitate more accurate interpretation of functional variability and help establish optimal adapter dosing ranges.

*In vivo* validation would also be essential to confirm tumor-specific killing, safety and potential immunogenicity due to repeated adapter injection. A key consideration for modular CAR systems such as SAM/AdCAR platforms is their dependence on repeated administration of adapter molecules to maintain therapeutic activity, which introduces pharmacokinetic constraints and potential immunogenicity risks. The biological activity of these systems is directly governed by the availability, distribution, and serum half-life of the adapter molecules, making efficacy highly sensitive to dosing schedule and clearance kinetics^46^. Rapidly cleared adapters allow tighter temporal control but require frequent re-dosing, whereas longer half-life formats increase exposure but may reduce controllability and increase toxicity risk^47^. Importantly, as with other biopharmaceuticals and CAR-based therapies, repeated exposure to engineered proteins may induce anti-drug immune responses that can reduce bioavailability, limit re-administration, and impair therapeutic efficacy over time^48^. Together, these pharmacokinetic and immunological constraints represent important translational considerations for adapter CAR strategies and highlight the need for systematic evaluation of dosing regimens and immunogenicity *in vivo*.

Beyond AdCAR, alternative AND-gate strategies based on other modular CAR systems have been recently investigated, notably the RevCAR system. RevCAR T cells are, similar to AdCAR T cells, activated via bifunctional adapter molecules that mediate target engagement. Instead of a chemically added tag like biotin, the RevCAR system conveys binding via peptide tags. A dual-RevCAR configuration distributes the stimulatory and co-stimulatory signaling domains across two separate CAR molecules, enabling combinatorial antigen sensing. This approach has shown encouraging preclinical results in colorectal cancer targeting CEA and EpCAM, a relevant setting for AND-gating due to their expression in some healthy tissues^49,50^. In this context, combinations such as CD318 with CD66c or TSPAN8 (or other PDAC targets) could be explored within RevCAR formats, potentially offering reduced manufacturing complexity, and donor-to-donor variability in this setting could be further evaluated.

Despite these challenges, our findings demonstrate that combinatorial AdCAR strategy provides a flexible and controllable framework for targeting heterogeneous solid tumors. By exploiting differences in antigen expression, this approach enables discrimination between tumor and healthy tissues, thereby reducing the risk of on-target/off-tumor toxicity. The modular nature of AdCAR allows rapid retargeting in response to antigen escape, while the reversibility of adapter binding offers an additional safety advantage, as CAR activity can be halted by withdrawal of the adapter. However, patient-specific dosing remains a critical consideration for clinical translation.

In conclusion, multi-antigen AdCAR platforms represent a promising and adaptable strategy for next-generation immunotherapy in PDAC. The AND-gate design enables potent and selective tumor killing while preserving T cell function and minimizing toxicity. Nevertheless, successful clinical translation will require careful optimization of target selection, dosing strategies, and patient-specific parameters to ensure both efficacy and safety.

## 5 Data Availability Statement

Publicly available data were retrieved from: GEPIA2^22^ (http://gepia2.cancer-pku.cn/#index) and EMBL-EBI/Expression Atlas^23^ (https://www.ebi.ac.uk/gxa/experiments/E-GTEX-8/.).

The datasets used and/or analyzed in this paper are available from the corresponding authors on reasonable request.

## 6 Ethics statement

PBMCs were isolated from the whole blood of healthy anonymous volunteers at Miltenyi Biotec, with written informed consent as approved by the local ethics committee of Ärztekammer Nordrhein (2020272). All blood samples were handled following the required ethical and safety procedures.

## 7 Author Contributions

C.D and D.S conceived the study and developed the experimental strategies. C.D performed the *in vitro* work and data analysis, as well as the manuscript preparation. K.V supported C.D for the cytotoxicity assay handling and manuscript revision. D.S and O.H contributed to supervision and manuscript revision.

## 9 Acknowledgments

The authors thank Ulrika Bader and Nele Knelangen for the anti-biotin AdCAR cloning and LV preparation. The authors further thank Nele Knelangen for the manuscript revision. The authors thank Lena Willnow and Niels Werchau for the biotinylation protocol. Finally, the authors thank Aleksej Frolov, Stefan Tomiuk and Michael Liu for their support with the bioinformatic tools.

The authors would like to acknowledge the use of BioRender.com with a valid license in the creation of figure 1.A (https://BioRender.com/xk506cn), figure 3.H (https://BioRender.com/r3vd1my), figure 4.F (https://BioRender.com/1ye01mn), figure 4.G (https://BioRender.com/ntvbcu5), supplementary figure 1 (https://BioRender.com/it2j7c4) and supplementary figure 2 (https://BioRender.com/18wpen0), all licensed under CC BY 4.0.

## 11 Generative AI statement

The author(s) declare that ChatGPT (GPT-5.3, OpenAI) was used in the creation of this manuscript, to improve the readability and language. After using this tool, the authors reviewed the content critically and thoroughly, edited it wherever needed, and take full responsibility for the content of the publication. All scientific content, interpretations, and conclusions were generated and verified by the authors.

## Supplementary figures

**Supplementary table 1.** List of Miltenyi antibodies used for flow cytometric analysis.

| Target | Clone | Conjugate |
| --- | --- | --- |
| CD318 | REA194, human | Unconjugated or PE |
| CD66c | REA414, human | Unconjugated or PE |
| TSPAN8 | REA443, human | Unconjugated or PE |
| CD19 | REA675, human | Unconjugated |
| AdCAR detection reagent | na | PE |
| Anti-biotin | REA746, human | PE |
| CD4 | REA623, human | VioGreen |
| CD69 | REA824, human | APC-Vio770 |
| CD137 (4-1BB) | REA765, human | APC |
| CD223 (LAG3) | REA351, human | VioBlue |
| CD279 (PD1) | REA1165, human | PE-Vio770 |
| 7-AAD | - | - |

**Supplementary figure 1.**
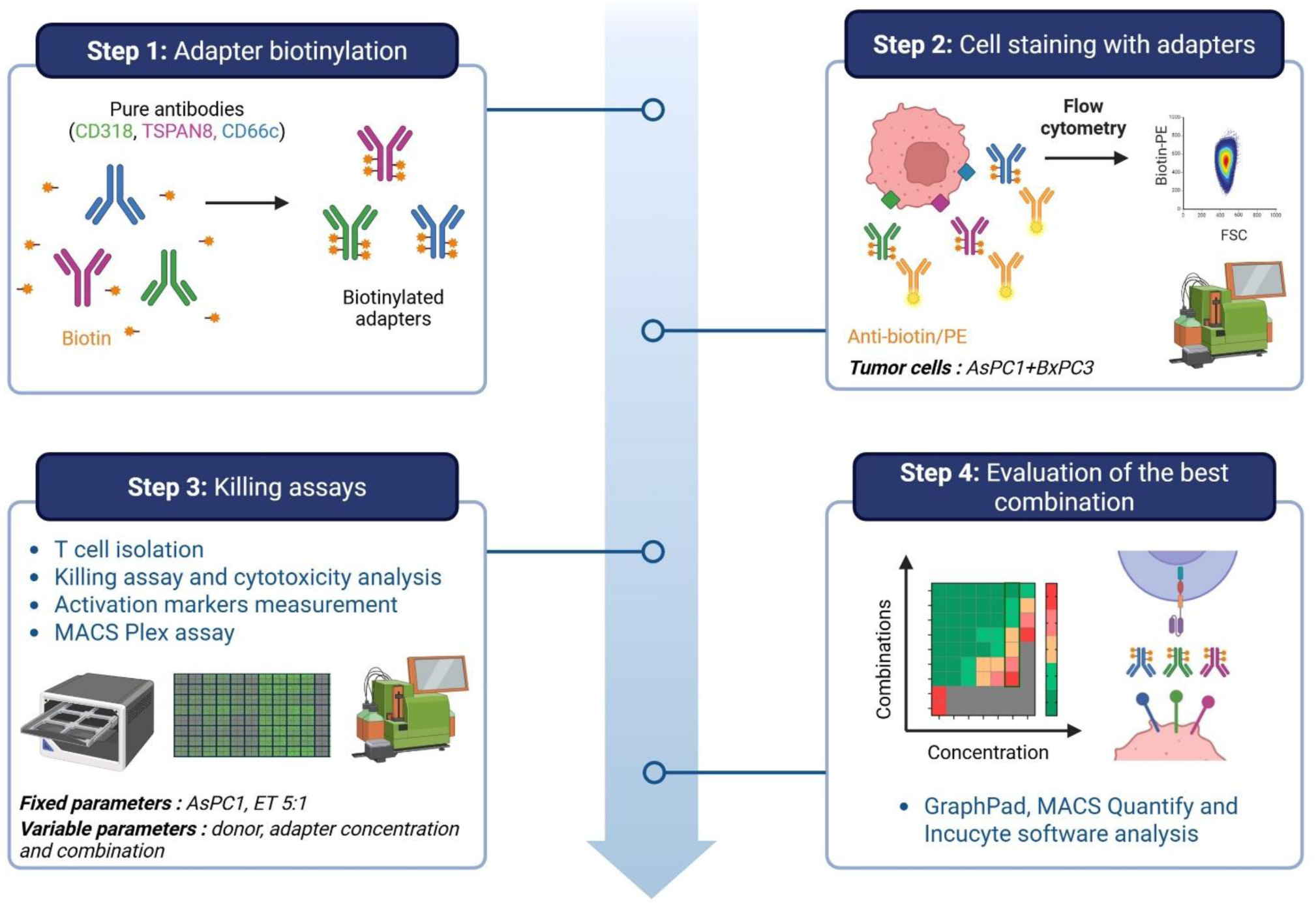
Complete workflow of the combinatorial AdCAR killing assay in vitro. Pure CD318, TSPAN8 and CD66c antibodies were first biotinylated, to make the adapters. A mixture of AsPC1 and BxPC3 (both PDAC cell lines) was stained and analyzed by flow cytometry to check the presence of the biotin tag. Adapters were then tested individually and in combination at different concentrations in a 4-day killing assay. T cells were isolated from different donors and co-cultured with AsPC1 cells at a 5:1 effector-to-target (E:T) ratio. Best combinations and the activation threshold were determined based on cytotoxicity abilities, activation markers and cytokines release. Data was processed using GraphPad and MACS Quantify to evaluate the best combinations. Created with BioRender.com.

**Supplementary figure 2.**
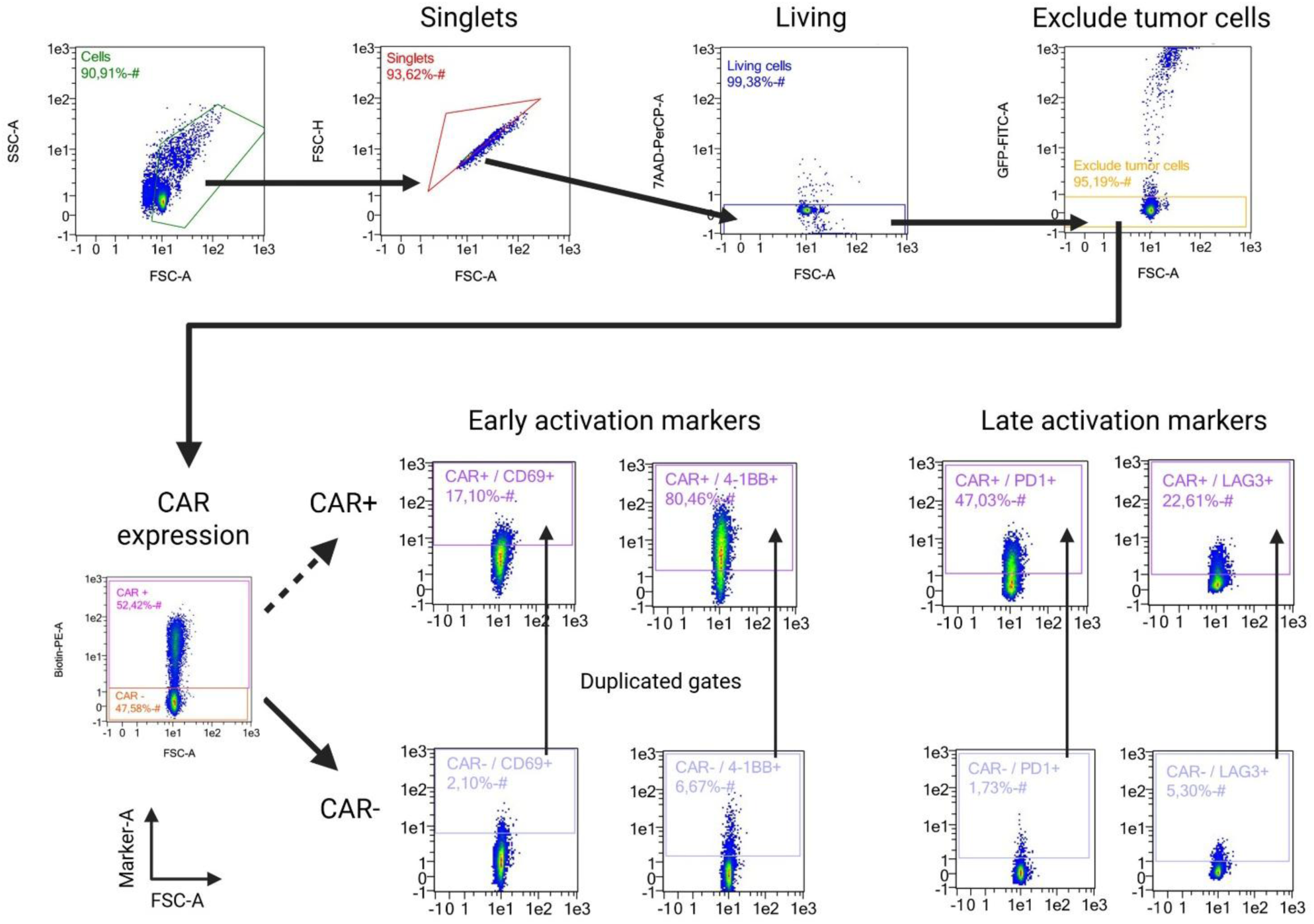
Gating strategy for activation marker analysis of immune cells and tumor cells co-cultures by flow cytometry. Initial gating steps excluded debris (FSC-A vs SSC-A), doublets (FSC-A vs FSC-H), dead cells (FSC-A vs 7AAD/PerCP-A) and tumor cells (FSC-A vs GFP/FITC-A). CAR⁺ T cells were gated as biotin-PE⁺. Early activation markers were assessed by the expression of CD69 and 4-1BB. Late activation markers were assessed by the expression of PD1 and LAG3. Activation-marker gates were established using CAR⁻ cells and the non-transduced control (NTC) and then applied to the CAR⁺ population to ensure that activation marker assessment was based on uniform, unbiased thresholds across all T cell subsets. Created with BioRender.com.

**Supplementary figure 3.**
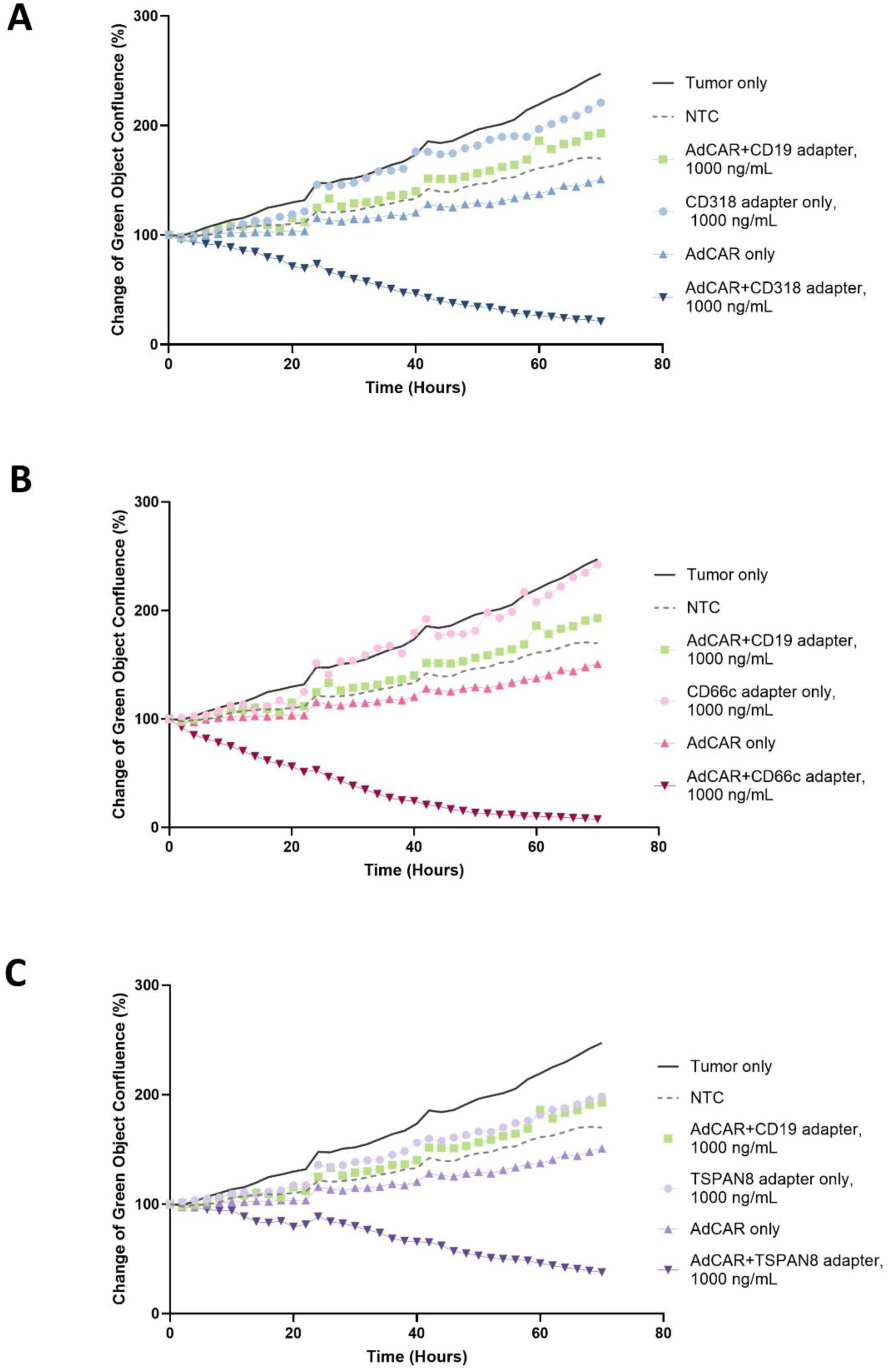
In vitro AdCAR killing assay using high concentrations of individual adapters and corresponding controls. Killing curves show GFP-positive tumor cell confluence (%) normalized to the first time point for AdCAR T cells combined with 1000 ng/mL of **A)** CD318, **B)** CD66c, or **C)** TSPAN8 adapter in co-culture with AsPC1 cells at an E:T ratio of 5:1. Control conditions included tumor cells alone, non-transduced T cells (NTC), adapter alone, AdCAR T cells without adapter, and AdCAR T cells combined with a CD19 adapter as a biotinylated isotype control (AsPC1 cells do not express CD19). Data points represent the mean of technical triplicates for each condition.

**Supplementary figure 4.**
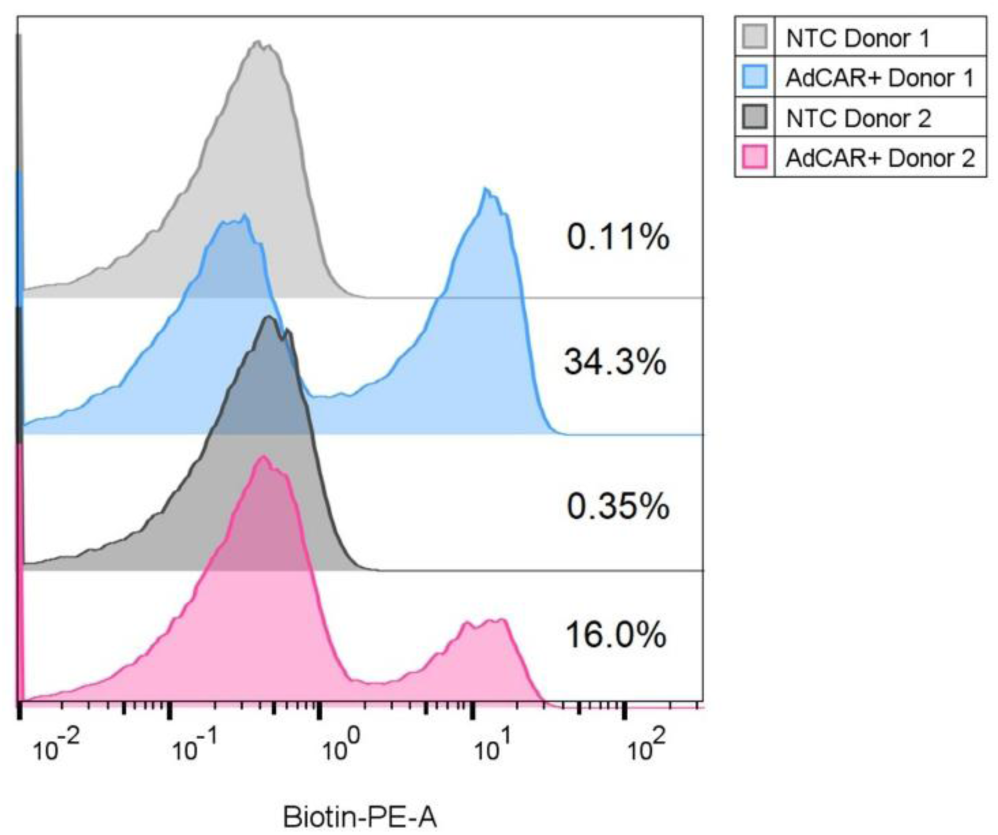
Density plots from flow cytometry showing AdCAR transduction efficiency on T cells from both donors, at day 10 after isolation. Samples were stained with AdCAR detection reagent PE-conjugated. Percentages indicate the positive fractions determined by using the negative control for gating.

**Supplementary figure 5.**
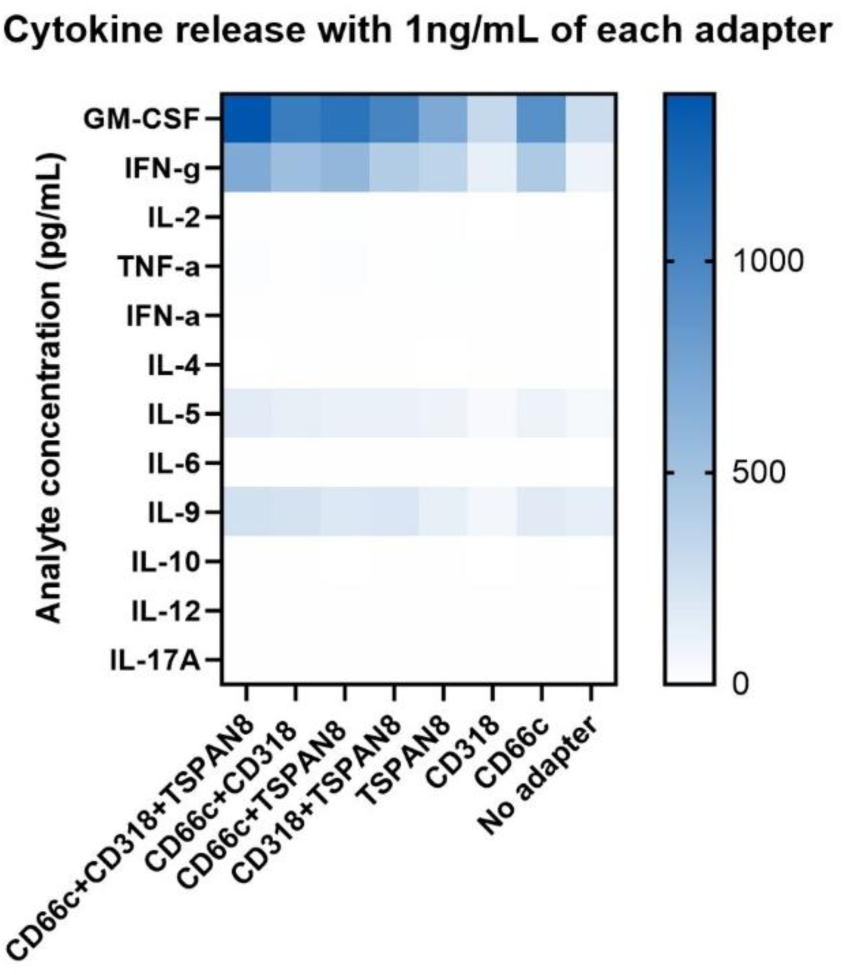
Heatmap summary of cytokine released at assay endpoint for each adapter combination at 1 ng/mL in donor 1. The analyte concentration is measured in pg/mL.

